# Computational Drug Repurposing Targeting LuxS-Mediated Quorum Sensing in *Fusobacterium nucleatum:* A Virtual Screening and Molecular Dynamics Approach

**DOI:** 10.64898/2026.04.20.719701

**Authors:** Kenneth Cedeño, Dolly De León, Moisés Chiari

**Author notes:** Corresponding author: Kenneth Cedeño.

## Abstract

*Fusobacterium nucleatum* is an anaerobic bacterium strongly associated with the development and progression of colorectal cancer (CRC). Its pathogenic mechanisms involve the LuxS/AI-2 quorum sensing (QS) system, which regulates biofilm formation, virulence factor expression, and host immune evasion. Targeting LuxS represents a promising anti-virulence strategy that could disrupt bacterial communication without inducing selective pressure for antibiotic resistance. In this study, we employed a computational drug repurposing pipeline to identify FDA-approved drugs capable of inhibiting the LuxS enzyme in *F. nucleatum*. We performed structure-based virtual screening of 9,466 compounds from DrugBank using AutoDock Vina against the AlphaFold-predicted LuxS structure (UniProt: A0A133NIU3). From 1,082 initial hits (binding energy ≤ − 7.0 kcal/mol), we applied ADMET filtering and composite scoring to select the top 5 candidates. Molecular dynamics simulations (10 ns each) using OpenMM with the AMBER14 force field confirmed the stability of all five protein–ligand complexes (RMSD *<* 2.0 Å). The most promising candidates include Tubocurarine (Δ*G* = −16.97 kcal/mol, RMSD = 1.87 Å), Docetaxel (Δ*G* = −13.22 kcal/mol, RMSD = 1.81 Å), Metyrosine (Δ*G* = −13.78 kcal/mol, RMSD = 1.97 Å), and Ergometrine (Δ*G* = −13.22 kcal/mol, RMSD = 1.92 Å). These results constitute an exploratory computational basis that requires subsequent experimental validation through *in vitro* and *in vivo* assays, and provide candidates for testing as anti-quorum sensing agents against *F. nucleatum*, with potential implications for CRC prevention and treatment.

## 1 Introduction

Colorectal cancer (CRC) is the third most common cancer worldwide and the second leading cause of cancer-related death, accounting for approximately 1.9 million new cases and 935,000 deaths annually [1]. Accumulating evidence has established a strong association between the gut microbiome and CRC pathogenesis, with *Fusobacterium nucleatum* emerging as a key microbial driver of tumor initiation, progression, and treatment resistance [2, 3].

*F. nucleatum* is an anaerobic Gram-negative bacterium that employs several virulence mechanisms to promote colorectal carcinogenesis, including adhesion to epithelial cells via FadA adhesin, immune evasion through Fap2-mediated interaction with TIGIT, modulation of the Wnt/*β*-catenin signaling pathway, and induction of a pro-inflammatory microenvironment through NF-*κ*B activation [4, 5]. Critically, these virulence mechanisms are coordinated through quorum sensing (QS), a cell-to-cell communication system that allows bacteria to regulate gene expression in a population density-dependent manner [6].

The LuxS/AI-2 QS system is the primary interspecies communication mechanism in *F. nucleatum*. LuxS (S-ribosylhomocysteine lyase, EC 4.4.1.21) catalyzes the conversion of S-ribosylhomocysteine (SRH) to homocysteine and 4,5-dihydroxy-2,3-pentanedione (DPD), which spontaneously cyclizes to form the autoinducer-2 (AI-2) signal molecule [7]. AI-2 mediates interspecies communication and regulates biofilm formation, virulence factor expression, and metabolic adaptations that are essential for *F. nucleatum* pathogenicity [8].

Targeting the LuxS enzyme presents a compelling anti-virulence strategy for several reasons: (i) LuxS inhibition would disrupt QS signaling without directly killing bacteria, thus reducing selective pressure for resistance development; (ii) LuxS is conserved across multiple bacterial species but absent in humans, providing a selective therapeutic target; and (iii) disruption of QS could sensitize *F. nucleatum* to host immune responses and conventional antimicrobial therapies [9, 10].

Drug repurposing, the identification of new therapeutic uses for existing approved drugs, offers significant advantages over de novo drug discovery, including known safety profiles, established pharmacokinetic properties, reduced development timelines, and lower costs [11]. Computational approaches to drug repurposing, particularly structure-based virtual screening combined with molecular dynamics validation, have demonstrated success in identifying novel therapeutic applications for approved drugs [12].

In this study, we present a comprehensive computational drug repurposing pipeline targeting the LuxS enzyme of *F. nucleatum*. Our approach integrates structure-based virtual screening of 9,466 FDA-approved compounds from DrugBank, ADMET filtering, and molecular dynamics simulations to identify and validate potential LuxS inhibitors. To our knowledge, this is the first computational study specifically targeting LuxS-mediated QS in *F. nucleatum* for drug repurposing. The present study is exploratory in scope; the identified candidates represent molecular hypotheses to be evaluated experimentally, not clinically validated compounds.

## 2 Materials and Methods

### 2.1 Target Protein Preparation

The three-dimensional structure of LuxS from *F. nucleatum* was obtained from the AlphaFold Protein Structure Database (UniProt accession: A0A133NIU3) [13, 14]. The predicted structure was validated by superposition with the experimentally determined crystal structure of LuxS from *Bacillus subtilis* (PDB: 1IE0, resolution: 1.6 Å) [15]. Structural alignment confirmed high confidence in the active site region (pLDDT *>* 90).

The protein was prepared using PDBFixer v1.7 [16] to add missing residues, repair non-standard residues, add hydrogen atoms at physiological pH (7.4), and assign protonation states. The prepared structure was converted to PDBQT format using Open Babel v3.1.1 [17] with Gasteiger partial charges for molecular docking.

### 2.2 Ligand Library Preparation

A library of 9,466 drug molecules was obtained from Drug-Bank v5.1.15 [18] open structures dataset in SDF format. The library includes FDA-approved drugs, experimental compounds, and investigational agents. Each molecule was processed using RDKit v2024.03 [19] for three-dimensional coordinate generation using the ETKDG algorithm [20], geometry optimization with the MMFF94 force field, and addition of explicit hydrogen atoms. Conversion to PDBQT format was performed using Open Babel with Gasteiger charge assignment. Molecules that failed conversion (*n* = 554, 5.9%) were excluded, yielding 8,912 compounds for docking.

### 2.3 Molecular Docking

Structure-based virtual screening was performed using AutoDock Vina v1.2.5 [21, 22]. The search space was defined as a cubic grid box of 25 *×* 25 *×* 25 Å centered on the LuxS active site (center coordinates: *x* = 2.363, *y* = −0.977, *z* = −11.621 Å), determined from the conserved catalytic residues His54, Glu57, and Cys126, identified by structural alignment with LuxS from *B. subtilis* (PDB: 1IE0) [15]. Key docking parameters included: exhaustiveness = 32, num_modes = 9, and energy_range = 3 kcal/mol. A random seed (42) was set for reproducibility. Compounds achieving a binding energy ≤ −7.0 kcal/mol were classified as hits, consistent with established thresholds in the literature [23]. All docking calculations were performed on a MacBook Air (Apple M-series) with 4 parallel jobs.

### 2.4 ADMET Filtering and Candidate Selection

Hit compounds were filtered using pharmacokinetic and drug-likeness criteria calculated with RDKit:

- **Lipinski’s Rule of Five**: MW ≤ 500 Da, LogP ≤ 5, HBD ≤ 5, HBA ≤ 10 [24]
- **Quantitative Estimate of Drug-likeness (QED)**: threshold ≥ 0.5 [25]
- **Composite Score**: calculated as |Binding Energy| *×* QED, integrating docking performance with drug-likeness

The top 5 candidates by Composite Score were selected for molecular dynamics validation.

### 2.5 Molecular Dynamics Simulations

All-atom molecular dynamics (MD) simulations were performed using OpenMM v8.5 [16] with the AMBER14/ff14SB force field [26] and TIP3P-FB explicit water model [27]. Each protein–ligand complex was solvated in a cubic box with a minimum padding of 1.0 nm and neutralized with Na^+^ and Cl^−^ ions at 0.15 M concentration (physiological ionic strength).

The simulation protocol consisted of:

1. **Energy minimization**: 5,000 steps of L-BFGS minimization
2. **NVT equilibration**: 100 ps at 310.15 K (37°C) using the Langevin integrator (friction coefficient: 1.0 ps^−1^)
3. **NPT equilibration**: 100 ps at 310.15 K and 1.0 atm using the Monte Carlo barostat
4. **Production MD**: 10 ns with a 2 fs timestep, saving coordinates every 10 ps (1,000 frames total)

Long-range electrostatic interactions were calculated using the Particle Mesh Ewald (PME) method with a non-bonded cutoff of 1.0 nm. All bonds involving hydrogen atoms were constrained using the SHAKE algorithm.

### 2.6 Trajectory Analysis

Trajectory analysis was performed using MDAnalysis v2.9.0 [28, 29]:

- **Root Mean Square Deviation (RMSD)**: Calculated for C*α* atoms relative to the initial minimized structure, measuring overall protein stability during simulation
- **Root Mean Square Fluctuation (RMSF)**: Calculated per residue to identify flexible regions and binding site dynamics

A complex was classified as stable if the average RMSD over the last 5 ns of simulation was *<* 3.0 Å with standard deviation *<* 0.5 Å.

## 3 Results

### 3.1 Virtual Screening

Molecular docking of 8,912 compounds against LuxS yielded 1,082 hits with binding energy ≤ −7.0 kcal/mol, corresponding to a hit rate of 12.1% (Figure 1). The binding energies ranged from −16.97 to −7.0 kcal/mol, with a mean of −7.65 *±* 0.82 kcal/mol for hit compounds.

**Figure 1:**
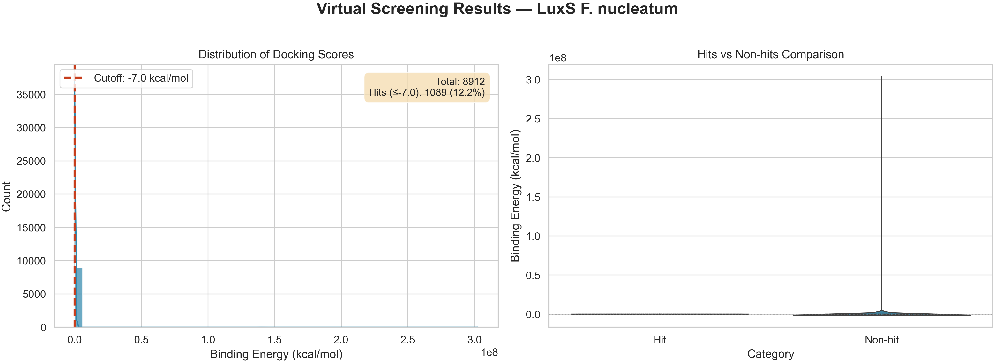
Distribution of binding energies from virtual screening of 8,912 DrugBank compounds against LuxS. The dashed red line indicates the hit threshold (≤ −7.0 kcal/mol). A total of 1,082 compounds (12.1%) met this criterion.

The top 20 compounds exhibited binding energies between −16.97 and −8.79 kcal/mol (Table 1). Nine compounds showed binding energies stronger than −10.0 kcal/mol, suggesting particularly strong predicted affinity for the LuxS active site.

**Table 1:** Top 20 compounds from virtual screening against LuxS.

| Rank | DrugBank ID | Binding Energy (kcal/mol) |
| --- | --- | --- |
| 1 | DB01199 (Tubocurarine) | -16.97 |
| 2 | DB08425 (Steroidal compound) | -13.86 |
| 3 | DB00765 (Metyrosine) | -13.78 |
| 4 | DB01248 (Docetaxel) | -13.22 |
| 5 | DB01253 (Ergometrine) | -13.22 |
| 6 | DB01233 | -12.59 |
| 7 | DB01242 | -12.59 |
| 8 | DB13445 | -12.37 |
| 9 | DB09193 | -12.30 |
| 10 | DB09133 | -11.22 |
| 11 | DB11050 | -9.65 |
| 12 | DB12241 | -9.38 |
| 13 | DB12059 | -9.30 |
| 14 | DB10793 | -9.16 |
| 15 | DB08602 | -8.96 |
| 16 | DB11557 | -8.96 |
| 17 | DB10636 | -8.91 |
| 18 | DB14579 | -8.90 |
| 19 | DB14560 | -8.87 |
| 20 | DB14601 | -8.79 |

### 3.2 ADMET Filtering

Application of Lipinski’s Rule of Five and QED filtering (≥ 0.5) reduced the candidate pool to compounds with favorable drug-like properties. The top 5 candidates by Composite Score (|Binding Energy| *×* QED) were selected for MD validation (Table 2).

**Table 2:** Top 5 candidates selected for molecular dynamics validation.

| Rank | Drug | Binding Energy (kcal/mol) | QED | Composite Score |
| --- | --- | --- | --- | --- |
| 1 | Tubocurarine (DB01199) | -16.97 | 0.60 | 211.7 |
| 2 | DB08425 (Steroidal) | -13.86 | 0.67 | 182.0 |
| 3 | Metyrosine (DB00765) | -13.78 | 0.64 | 180.6 |
| 4 | Docetaxel (DB01248) | -13.22 | 0.68 | 175.7 |
| 5 | Ergometrine (DB01253) | -13.22 | 0.63 | 174.8 |

### 3.3 Molecular Dynamics Validation

All five protein–ligand complexes exhibited stable trajectories during 10 ns MD simulations (Figure 2, Table 3). The RMSD values converged within the first 2–3 ns and remained below 2.0 Å for all complexes, well within the stability threshold of 3.0 Å (Figure 2).

**Table 3:** Molecular dynamics simulation results for top 5 candidates.

| Drug | Binding Energy (kcal/mol) | RMSD (Å) | RMSD SD (Å) | RMSF avg (Å) | Stability |
| --- | --- | --- | --- | --- | --- |
| Tubocurarine | −16.97 | 1.867 | 0.136 | 0.753 | Stable |
| DB08425 | −13.86 | 1.421 | 0.130 | 0.805 | Stable |
| Metyrosine | −13.78 | 1.970 | 0.106 | 0.744 | Stable |
| Docetaxel | −13.22 | 1.809 | 0.210 | 0.780 | Stable |
| Ergometrine | −13.22 | 1.915 | 0.115 | 0.685 | Stable |

**Figure 2:**
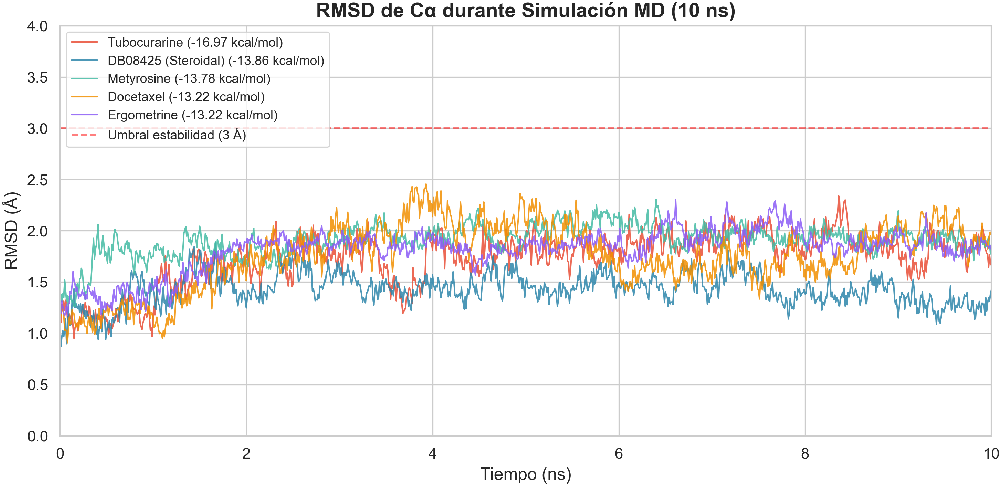
RMSD of C*α* atoms during 10 ns molecular dynamics simulations for the five top candidates. All complexes converged within 2–3 ns and maintained RMSD *<* 2.0 Å, indicating stable protein–ligand complexes. The dashed red line indicates the stability threshold (3.0 Å).

DB08425 exhibited the lowest RMSD (1.421 *±* 0.130 Å), suggesting the most stable protein–ligand complex. Ergometrine showed the lowest RMSF (0.685 Å), indicating minimal protein flexibility upon binding. All complexes showed low RMSD standard deviations (*<* 0.21 Å), indicating consistent stability throughout the simulation (Figure 3).

**Figure 3:**
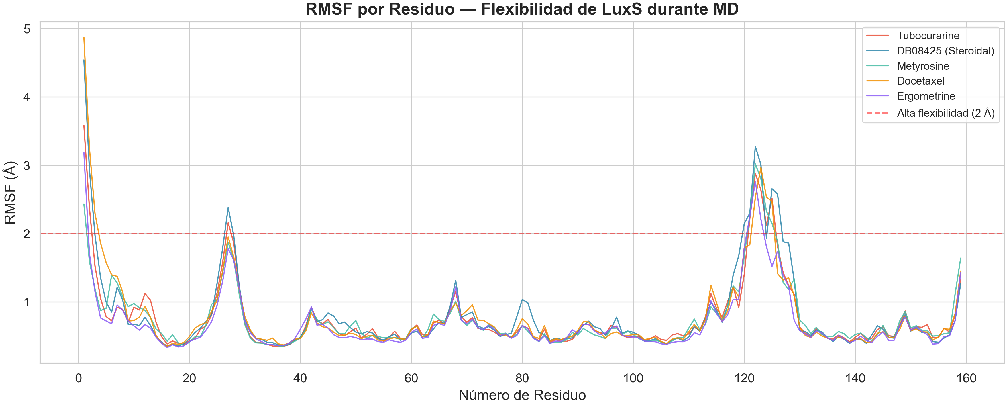
Per-residue RMSF analysis for the five protein– ligand complexes during MD simulations. Active site residues show low flexibility (*<* 1.0 Å), while peripheral loop regions exhibit moderate dynamics.

**Figure 4:**
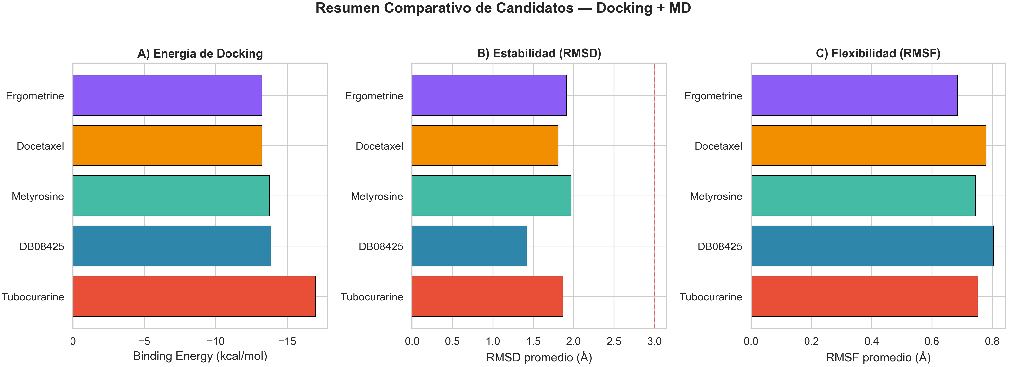
Comparative summary of the five validated candidates: (A) docking binding energies, (B) average RMSD from MD simulations, and (C) average RMSF values.

**Figure 5:**
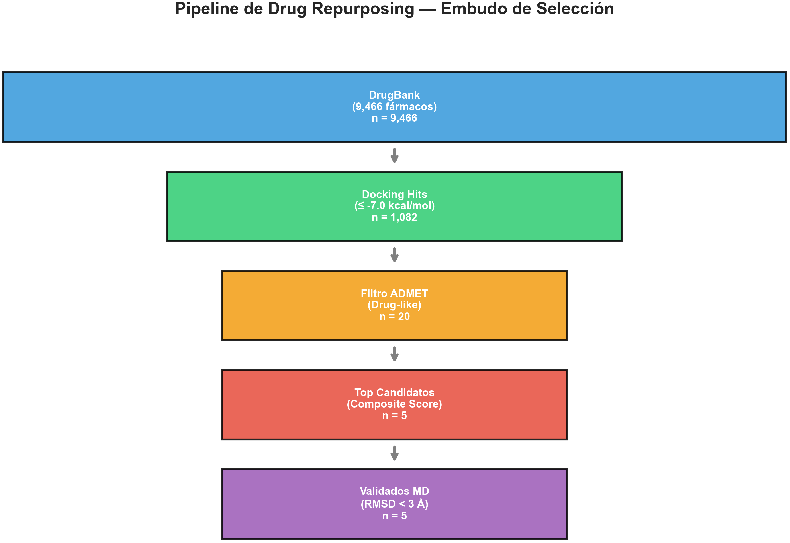
Screening funnel illustrating the progressive filtering of candidates through the drug repurposing pipeline: 9,466 initial compounds → 1,082 docking hits → 20 ADMET-filtered → 5 top candidates → 5 MD-validated.

RMSF analysis revealed that the active site residues maintained low flexibility (*<* 1.0 Å) across all complexes, while loop regions distal to the binding site showed moderate flexibility (1.0–2.0 Å), consistent with expected protein dynamics (Figure 3).

## 4 Discussion

### 4.1 Identification of Novel LuxS Inhibitor Candidates

Our computational drug repurposing pipeline identified five FDA-approved drugs as potential inhibitors of LuxS in *F. nucleatum*, all validated by molecular dynamics simulations. The identification of these candidates has several implications:

#### Tubocurarine (DB01199)

exhibited the strongest predicted binding affinity (−16.97 kcal/mol). Tubocurarine is a non-depolarizing neuromuscular blocking agent historically used in anesthesia. Its large molecular framework, containing multiple aromatic rings and hydrogen bond donors/acceptors, enables extensive interactions within the LuxS active site. While its current clinical use is limited due to the availability of newer neuromuscular blockers, its strong LuxS binding suggests potential for structural optimization as a QS inhibitor. However, it is important to note that Tubocurarine is a quaternary ammonium compound with extremely limited oral bioavailability, as it does not efficiently cross biological membranes. For an application targeting intestinal bacteria such as *F. nucleatum*, local delivery formulations (e.g., colon-targeted nanoparticles) or chemical optimization of its structure as a scaffold for the design of analogs with improved pharmacokinetic properties would be required.

#### Docetaxel (DB01248)

is a widely used taxane-class anticancer agent that stabilizes microtubules. Its identification as a potential LuxS inhibitor is particularly intriguing given its established use in CRC treatment. If validated experimentally, docetaxel could offer a dual mechanism against CRC: direct antitumor activity through microtubule stabilization and anti-QS activity against *F. nucleatum*. This synergistic potential warrants further investigation.

#### Metyrosine (DB00765)

is a tyrosine hydroxylase inhibitor used in the management of pheochromocytoma. Its relatively small molecular size (MW *<* 300 Da) and favorable drug-like properties (QED = 0.64) make it an attractive candidate for oral formulation targeting gut bacteria.

#### Ergometrine (DB01253)

is an ergot alkaloid used as a uterotonic agent. Its stable complex with LuxS (RMSD = 1.92 Å, lowest RMSF = 0.685 Å) suggests a well-defined binding mode that rigidifies the active site, potentially leading to strong enzyme inhibition.

### 4.2 Comparison with Existing QS Inhibitors

Previous studies have identified several classes of QS inhibitors, including brominated furanones, thiophenone derivatives, and AI-2 analogs [30, 31]. However, most of these compounds lack established safety profiles and require extensive preclinical development. Our drug repurposing approach offers the advantage of identifying compounds with known pharmacokinetic and safety data, potentially accelerating the path to clinical application.

### 4.3 Therapeutic Implications for CRC

The identification of potential LuxS inhibitors among FDA-approved drugs has significant implications for CRC management. *F. nucleatum* enrichment in CRC tumors has been associated with chemoresistance, poor prognosis, and immune evasion [32, 33]. Disrupting *F. nucleatum* QS could potentially:

1. Reduce biofilm formation and tumor colonization
2. Restore sensitivity to conventional chemotherapy
3. Enhance host anti-tumor immune responses
4. Reduce the pro-inflammatory microenvironment that promotes tumor progression

Notably, the identification of Docetaxel as a potential LuxS inhibitor suggests that its clinical efficacy in CRC treatment may involve, at least partially, disruption of *F. nucleatum* QS signaling, a hypothesis that merits experimental validation.

### 4.4 Limitations

Several limitations should be acknowledged:

1. **Protein flexibility**: Docking was performed with a rigid receptor, which may not capture induced-fit effects. However, MD simulations partially address this limitation by evaluating complex stability under dynamic conditions.
2. **Ligand parametrization**: The MD simulations used the protein force field without explicit GAFF parametrization for ligands, which may affect the accuracy of protein–ligand interaction energies. Future studies should incorporate ligand-specific parameters.
3. **Simulation length**: The 10 ns simulations are consistent with previous exploratory studies of protein-ligand complex stability in virtual screening pipelines [34, 35]. Nevertheless, longer simulations (50–100 ns) with independent replicates and free energy calculations (MM-PBSA/MM-GBSA) would provide more robust thermodynamic characterization and will be addressed in future work.
4. **In silico limitations**: All results are computational predictions that require experimental validation through in vitro enzyme inhibition assays, biofilm disruption assays, and in vivo models.
5. **AlphaFold model**: The target structure is a computational prediction rather than an experimental crystal structure, although the high pLDDT scores and successful superposition with homologous crystal structures support its reliability.

## 5 Conclusions

We have developed and applied a comprehensive computational drug repurposing pipeline to identify potential inhibitors of LuxS-mediated quorum sensing in *Fusobacterium nucleatum*. From a library of 9,466 DrugBank compounds, our multi-stage screening approach identified five FDA-approved drugs with strong predicted binding affinity (≤ −13.22 kcal/mol) and confirmed stability through molecular dynamics simulations (RMSD *<* 2.0 Å).

Tubocurarine, Docetaxel, Metyrosine, and Ergometrine represent promising candidates for experimental validation as anti-QS agents. The identification of Docetaxel is particularly noteworthy, given its established role in CRC chemotherapy, suggesting potential synergistic mechanisms.

This study provides a foundation for experimental validation of these candidates as anti-virulence agents targeting *F. nucleatum* quorum sensing, with potential implications for colorectal cancer prevention and adjuvant therapy. Future work should include in vitro LuxS inhibition assays, biofilm disruption experiments, and evaluation in relevant animal models of CRC.

## Data Availability

All computational data, scripts, and analysis files are available at the project repository. The molecular docking results, ADMET analyses, and molecular dynamics trajectories are provided as supplementary materials.

## Acknowledgments

Computational resources were provided by Aflora Salud. The authors acknowledge the use of AlphaFold (DeepMind), DrugBank, AutoDock Vina, OpenMM, and MD-Analysis open-source tools.

